# Functional opsin patterning for *Drosophila* color vision is established through signaling pathways in adjacent object-detection neurons

**DOI:** 10.1101/2023.09.29.560118

**Authors:** Manabu Kitamata, Yoshiaki Otake, Hideaki Kitagori, Xuanshuo Zhang, Yusuke Maki, Rika Boku, Masato Takeuchi, Hideki Nakagoshi

## Abstract

Vision is mainly based on two different tasks, object detection and color discrimination through activities of photoreceptor (PR) cells. *Drosophila* compound eye consists of ∼800 ommatidia. Every ommatidium contains eight PR cells; six outer cells (R1-R6) and two inner cells (R7 and R8) by which object detection and color vision are achieved, respectively. Expression of opsin genes in R7 and R8 is highly coordinated through the instructive signal from R7 to R8, and two major ommatidial subtypes are distributed stochastically; pale type expresses Rh3/Rh5, while yellow type expresses Rh4/Rh6 in R7/R8. The homeodomain protein Defective proventriculus (Dve) is expressed in yellow-type R7 and in six outer PRs, and it is involved in Rh3 repression to specify the yellow-type R7. *dve* mutant eyes exhibited atypical coupling, Rh3/Rh6 and Rh4/Rh5, indicating that the Dve activity is required for proper opsin coupling. Surprisingly, Dve activity in R1 is required for instructive signal, whereas those in R6 and R7 block the signal. Our results indicate that functional coupling of two different neurons is established through signaling pathways from adjacent neurons that are functionally different.

**Summary Statement:** Dve activity is required for proper opsin coupling of two neurons that is established through signaling pathways from adjacent neurons that are functionally different.

## Introduction

In vertebrates, rod cells express *rhodopsin* (*rh*) and are involved in object detection in dim light, while cone cells, for example in humans, express one of three types of cone opsin, which absorb short (S, blue), medium (M, green), and long (L, red) wavelengths, respectively. Retinas have mosaic distribution of these cone cells, and color vision is achieved by comparing the outputs of PR cells that have different spectral sensitivities (Nathans, 1999).

*Drosophila* compound eye consists of ∼800 ommatidia. Every ommatidium contains eight PR cells: six outer cells (R1-R6) and two inner cells (R7 and R8). Outer PR cells express Rhodopsin1 (Rh1) and are involved in object (motion) detection, while color vision is achieved by inner PR cells, which express UV-sensitive opsins (Rh3 and Rh4) in R7, and blue- and green-sensitive opsins (Rh5 and Rh6) in R8, respectively. Expression of opsin genes in R7 and R8 is highly coordinated through the instructive signal from R7 to R8 (Chou et al., 1999), and two major subtypes of ommatidium are distributed stochastically; pale type (∼30%) expresses Rh3/Rh5, while yellow type (∼70%) expresses Rh4/Rh6 in R7/R8 (Mollereau and Domingos, 2005; Wernet and Desplan, 2004). Photo-sensitive structure rhabdomeres of R7 and R8 are vertically aligned on the same axis in an ommatidium. Thus, this arrangement with functional Rh coupling (Rh3/Rh5 and Rh4/Rh6) is thought to be critical for color vision, namely response to different wavelength.

PR differentiation is regulated by two steps: (1) cell-fate determination and axonal projection during larval development, and (2) terminal differentiation such as rhabdomere morphogenesis and opsin gene expression during pupal development (Mollereau et al., 2001). The early event of terminal differentiation is the specification of inner and R7 identities by *spalt* and *prospero* (*pros*) (Cook et al., 2003; Mollereau et al., 2001). Subsequent specification of R7 is performed by *orthodenticle* (*otd*) and *spineless* (*ss*) (Tahayato et al., 2003; Wernet et al., 2006). Expression of opsins in R7 and R8 is highly coordinated, and two major subtypes of ommatidium, pale and yellow, are established through the instructive signal from R7 to R8 within an ommatidium (Chou et al., 1999).

In response to R7 subtypes, transmission or blockade of the instructive signal is selected, and a bistable loop between *warts* (*wts*) and *melted* (*melt*) determines the state of Rh5 or Rh6 expression in R8 (Mikeladze-Dvali et al., 2005). Wts is a Ser/Thr kinase that is a core component of the Hippo signaling pathway involved in growth suppression. During ommatidial development, *wts* is necessary and sufficient for R8 to adopt the yellow-type identity, while *melt* plays the opposite role and induces the pale-type identity in R8. These two genes repress each other’s transcription to form a bistable loop (Anderson et al., 2017; Jukam and Desplan, 2011; Jukam et al., 2013; Thanawala et al., 2013). In addition, Activin and BMP signaling are required upstream of the Hippo pathway to establish expression of Rh5 or Rh6 in R8 (Wells et al., 2017). Involvement of Epidermal growth factor receptor, Rhomboid, and Hibris is also reported (Birkholz et al., 2009a; Birkholz et al., 2009b; Tan et al., 2020). However, mechanisms by which the instructive signal is transmitted from R7 to R8 remains unknown. Here, we provide evidence that transmission of the instructive signal is regulated by the activities of Defective proventriculus (Dve) in the outer PR cells, R1 and R6.

## Results and Discussion

### Abnormal Rhodopsin coupling in *dve* mutant ommatidia

The homeodomain transcription factor Defective proventriculus (Dve) is involved in various functions including cell-type specification, functional differentiation, and cell survival. Dve is expressed in all outer PR cells and in yellow-type R7 (yR7). Dve expression in yR7 depends on the activity of Ss and represses the pale-type opsin Rh3 to specify the yellow-type identity. In *dve* mutant eyes, Rh3 and Rh4 are coexpressed in yR7 due to derepression of Rh3 (Johnston et al., 2011). In *dve*^*1*^ mutant eyes, the ratio of Rh5/Rh6 in the R8 layer was nearly normal, however, their coupling to R7 Rhodopsins was quite abnormal (Fig. 1A-E). Rh3/Rh6 coupling in R7/R8 is rarely observed in wild-type ommatidia, and this coupling is thought to be a default state. In *dve*^*1*^ heterozygous control eyes, the Rh3/Rh6 coupling was observed only at 0.3%, whereas it was frequently observed at 23.1% in *dve*^*1*^ homozygous mutant eyes (Fig. 1A-C, and E). In addition, atypical coupling Rh3+Rh4/Rh5 was also observed at 14.8% (Fig. 1A, B, D, and E). This atypical coupling is never observed in wild-type ommatidia. Thus, transmission of the instructive signal from R7 to R8 appears to be randomized in *dve*^*1*^ mutant ommatidia. Because *dve*^*1*^ is a severe loss of function allele, we checked the effect of null allele *dve*^*L186*^. Surprisingly, a considerable amount of *dve*^*L186*^ mutant ommatidia (32.6%) exhibited atypical Rh3/Rh6 coupling, and almost all cells in R8 expressed Rh6 (Fig. 1F). In the absence of Dve activity, all R7 express Rh3 due to derepression of Rh3 in yR7. Thus, almost all ommatidia express Rh3 and Rh6 as a default state in *dve*^*L186*^ mutant eyes, indicating that the Dve activity is crucial for the instructive signal from R7 to R8.

**Figure 1.**
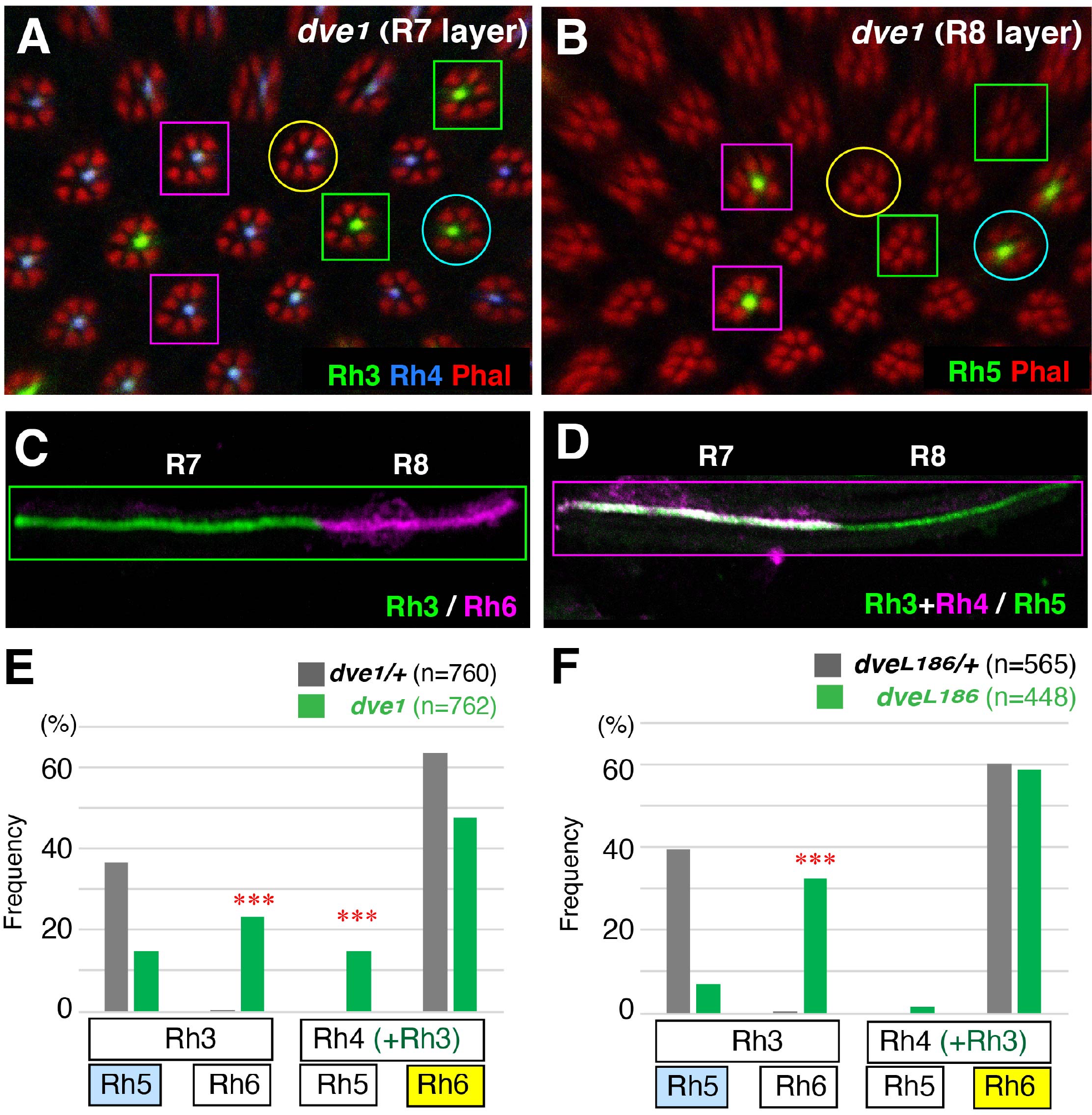
Rhodopsin coupling is abnormal in *dve* mutant ommatidia. (A-D) Rhodopsin expression in *dve1* mutant eyes (y*w eyflp/Y; FRT42D dve 1/FRT42D w+M*). (A, B) R7 and R8 layers of a *dve1* compound eye. Two typical types of ommatidia are outlined with circles (pale and yellow, Rh3/Rh5 and Rh4/Rh6, respectively). Atypical types of ommatidia are outlined with squares (green and magenta, Rh3/Rh6 and Rh3+Rh4/Rh5, respectively). Rhabdomeres are labeled with Phalloidin (red). Rh3 and Rh5 (green). Rh4 (blue). (C, D) Dissociated ommatidia show atypical Rhodopsin coupling as outlined with squares in A and B. Rh3 and Rh5 (green). Rh4 and Rh6 (magenta). (E) Rhodopsin coupling of *dve1/+* (*yw/Y; FRT42D dve1/+*, N=10 compound eyes, n=760 ommatidia) and *dve1/dve1* ommatidia (y*w eyflp/Y; FRT42D dve 1/ FRT42D w+M*, N=12, n=762) (F) Rhodopsin coupling of *dveL186/+* (*yw/Y; FRT42D dveL186/+*, N=7, n=565) and *dveL186/dveL186* ommatidia (y*w eyflp/Y; FRT42D dveL186/FRT42D w+M*, N=7, n=448). Atypical couplings in *dve* mutant ommatidia are indicated by asterisks (chi-square test, p<0.0001).

### Dve activity in outer PRs regulates proper coupling

To further examine functions of Dve for the instructive signal, *dve* mutation was introduced into the *ss* mutant background (*dve ss* double mutants). In *ss* mutant eyes, almost all ommatidia becomes to pale-type coupling (Rh3/Rh5) (Fig. 2A, C, and E). Heterozygous or homozygous *dve* mutation in the *ss* mutant background did not affect Rh3 expression in the R7 layer (Fig. 2A, B), whereas *dve ss* double mutant eyes significantly increased the number of Rh6-expressing R8 (Fig. 2D, F). These results further support the above notion that the Dve activity is essential for transmission of the instructive signal from R7 to R8. If the instructive signal is directly transmitted from pR7 to R8, the signal transduction between rhabdomeres is probable, because their cell bodies are separated by the outer PR, R1. Rhabdomere is a specialized structure where Rhodospins and actin are densely accumulated, and Rh3 itself might act as an instructive signaling molecule. It should be noted that, in *dve*^*L186*^ mutant ommatidia, the instructive signal is almost completely blocked despite Rh3 expression in all R7s.

**Figure 2.**
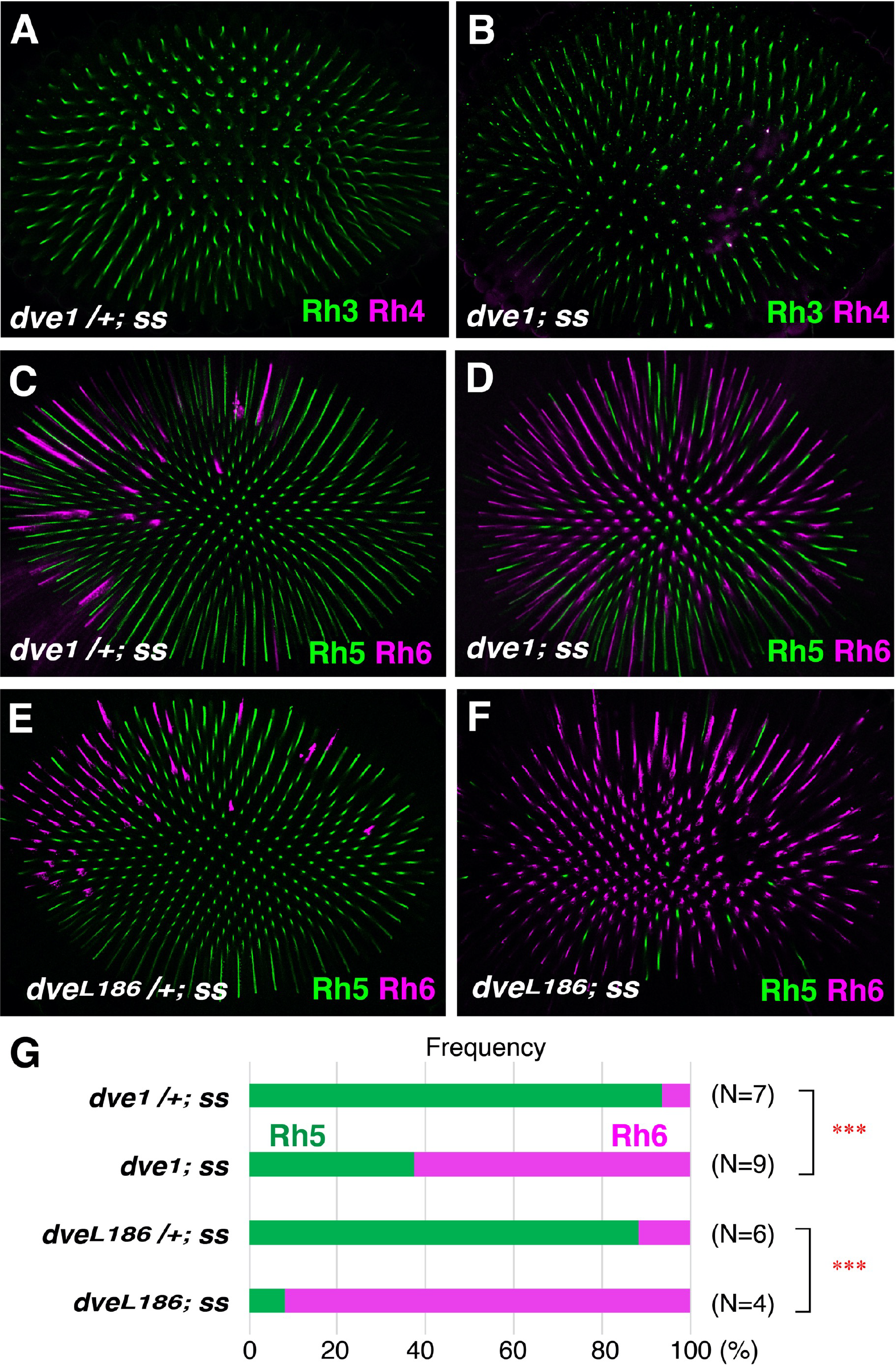
Dve activity in outer PRs regulates proper coupling. (A-F) Rhodopsin expression in *ss* mutant eyes [*dve1/+; ssD115*.*7* (A, C) and *dveL186/+; ssD115*.*7* (E)] and *dve ss* double mutant eyes [*dve1; ssD115*.*7* (B, D) and *dveL186; ssD115*.*7* (F)]. (A, B) Expression of Rh3 (green) and Rh4 (magenta) in the R7 layer. (C-F) Expression of Rh5 (green) and Rh6 (magenta) in the R8 layer. (G) Expression ratio of the R8 Rhodopsin. *dve1/+; ssD115*.*7* (N=7, n=2929), *dve1; ssD115*.*7* (N=9, n=2242), *dveL186/+; ssD115*.*7* (N=6, n=2515), and *dve1; ssD115*.*7* (N=4, n=1030). Rh6 is greatly increased in the *dve ss* double mutant background (Fischer’s exact test, p<0.0001).

Because Dve expression is undetectable in the pale-type R7 (pR7), it seems unlikely that the undetectable level of Dve expression in pR7 is critically required for sending the instructive signal from pR7. Therefore, we favor another possibility that the Dve activity in outer PRs is required for transmission of the instructive signal, because Dve is strongly expressed in outer PRs, R1-R6.

### Dve activity in R1 is required for instructive signaling

Rhabdomeres of R7 and R8 are vertically aligned on the same axis but their cell bodies are separated by the outer PR, R1. Based on the topological arrangement of their cell bodies, we hypothesized that the instructive signal is transmitted from R7 to R1, and then R1 to R8. To induce *dve* mutation in specific cell types, we used *GMR-flp* that expresses Flp recombinase after second mitotic wave, namely in R1, R6, and R7. In the MARCM system, *dve* mutant cells were labeled as GFP-expressing cells, and a mutant cell completely lost Dve expression. By using this mosaic system, Rhodopsin coupling was scored in cell-specific *dve* mutant ommatidia (Fig. 3). In R1 *dve* mutant ommatidia, only two types of Rh coupling were observed. These are the default state (Rh3/Rh6) and the yellow type (Rh4/Rh6), and Rh5 was never induced at all. This result clearly shows that the Dve activity in R1 is critically required for transmission of the instructive signal (Fig. 3A).

**Figure 3.**
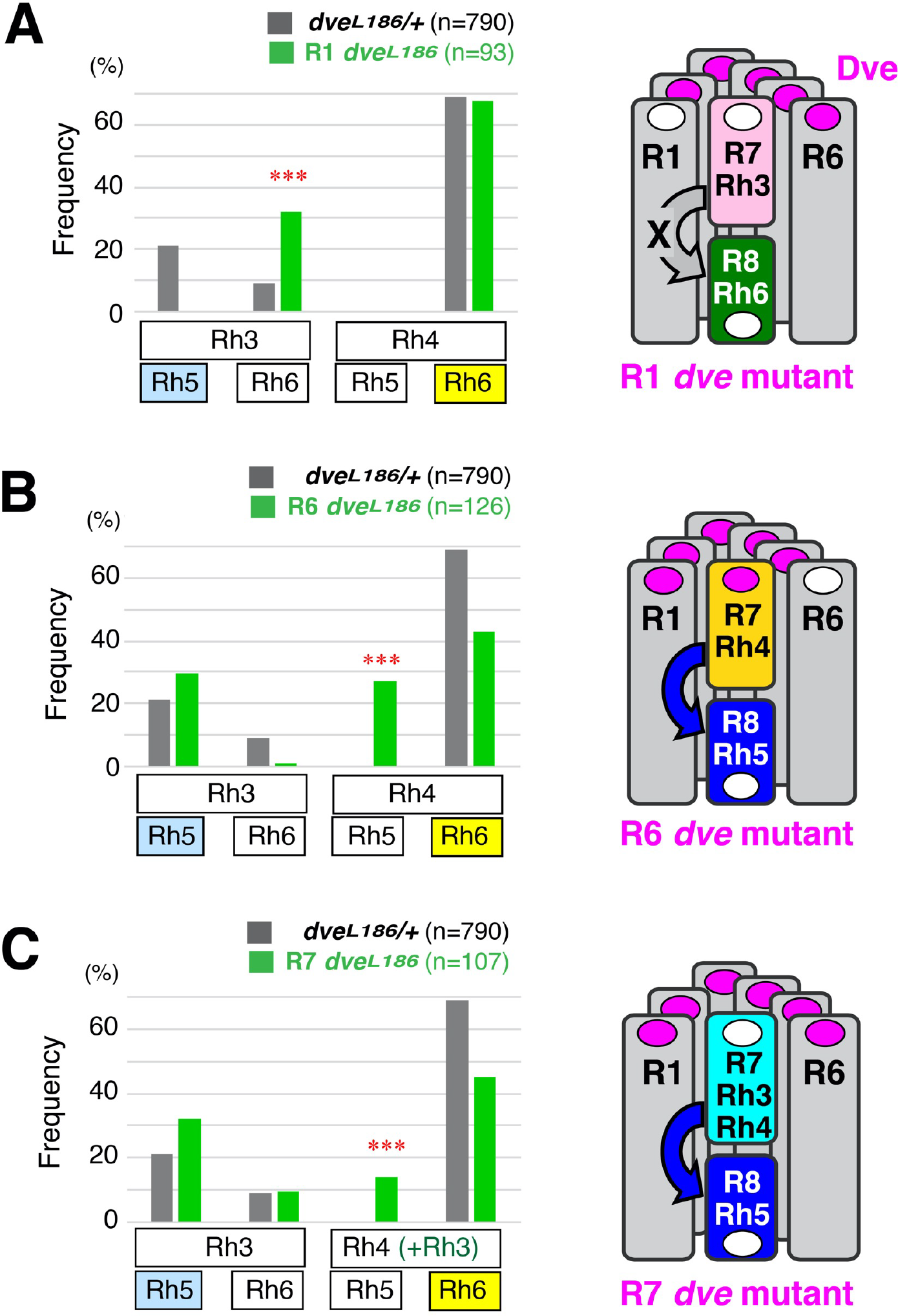
Two opposite functions of Dve in Rhodopsin coupling. Cell-specific *dve* mutations are induced with the MARCM system. (A) *dve* mutation in R1 (n=93). Atypical coupling (Rh3/Rh6) is indicated by asterisks (chi-square test, p<0.0001). (B) *dve* mutation in R6 (n=126). Atypical coupling (Rh4/Rh5) is indicated by asterisks (chi-square test, p<0.0001). (C) *dve* mutation in R7 (n=107). Atypical coupling (Rh3+Rh4/Rh5) is indicated by asterisks (chi-square test, p<0.0001). Atypical couplings with cell-specific *dve* mutations are schematically shown in right sides. Rectangles and circles represent rhabdomeres and nuclei of R cells, respectively. Dve expression is shown with magenta circles.

### Dve activities in R6 and R7 block the instructive signal

In R6 or R7 *dve* mutant ommatidia, atypical coupling Rh4/Rh5 was frequently observed (Fig. 3B, C). This coupling was also observed in *dve*^*1*^ mutant eyes (Fig. 1D) and reflects ectopic transmission of the instructive signal from Rh4-expressing yR7 to R8. This might be due to ectopic Rh3 expression in yR7. For instance, *rh3* knock down (KD) increased the number of default state ommatidia (Rh3/Rh6) to some extent. However, this contribution is only redundant, because ectopic transmission was frequently observed with *rh3* KD conditions in R7 *dve* mutants. Furthermore, R6 *dve* mutant ommatidia did not induce ectopic Rh3 expression in yR7, whereas they induced atypical coupling Rh4/Rh5 at 42.1% (Fig. 3B). Taken together, ectopic instructive signal can be generated in the absence of Dve activities in R6 or R7 rather than the ectopic Rh3 expression in R7.

### Mechanisms of Rhodopsin coupling

In human retinas, the orphan nuclear receptor NR2E3 (also known as PNR) is expressed in rod cells (Bumsted O’Brien et al., 2004; Cheng et al., 2004) and activates the expression of rod genes but represses that of cone genes (Peng et al., 2005). Interestingly, mutations in the NR2E3 gene not only cause defects in rod system but also change the sensitivity of cone cells. Their retinas are hypersensitive to blue light (S-cone) and have reduced sensitivity to green and red light (M- and L-cones), leading to ‘enhanced S-cone syndrome’ (Haider et al., 2000). The NR2E3 mutant phenotype is similar to that of our observation, because *dve* mutation in R1 or R6 affects Rhodopsin expression in R8. Although loss of function in rod cells affects functions in cone cells, it is due to misdifferentiation of rod precursor cells into the default cell type, S-cones, but not through abnormal intercellular signaling (Cheng et al., 2011).

Our results provide first evidence that functional opsin patterning (in R7/R8) for *Drosophila* color vision is established through signaling pathways in adjacent object-detection neurons (R1 and R6). Our model is shown in Fig. 4. In a default state, Rh5 expression is repressed by the Hippo pathway in R8. In pale-type ommatidia, Rh3 expression in R7 sends an instructive signal and relieves repression of Rh5 in R8 through disruption of the Hippo signaling pathway (Mikeladze-Dvali et al., 2005). The instructive signal from R7 activates Dve in adjacent R1, and that the activated Dve (Dve*) disrupts the Hippo signaling pathway in adjacent R8. In yellow-type ommatidia, Dve activities in R7 and R6 appear to repress the activation of Dve in R1, resulting in active Hippo signaling and Rh6 expression in R8. Additional mechanism is also possible that BMP and Activin are secreted from pale R7 to the surface of adjacent R1, and that they are actively transferred to their receptors in R8. This transfer mechanism might be regulated by the Dve activity in R1.

**Figure 4.**
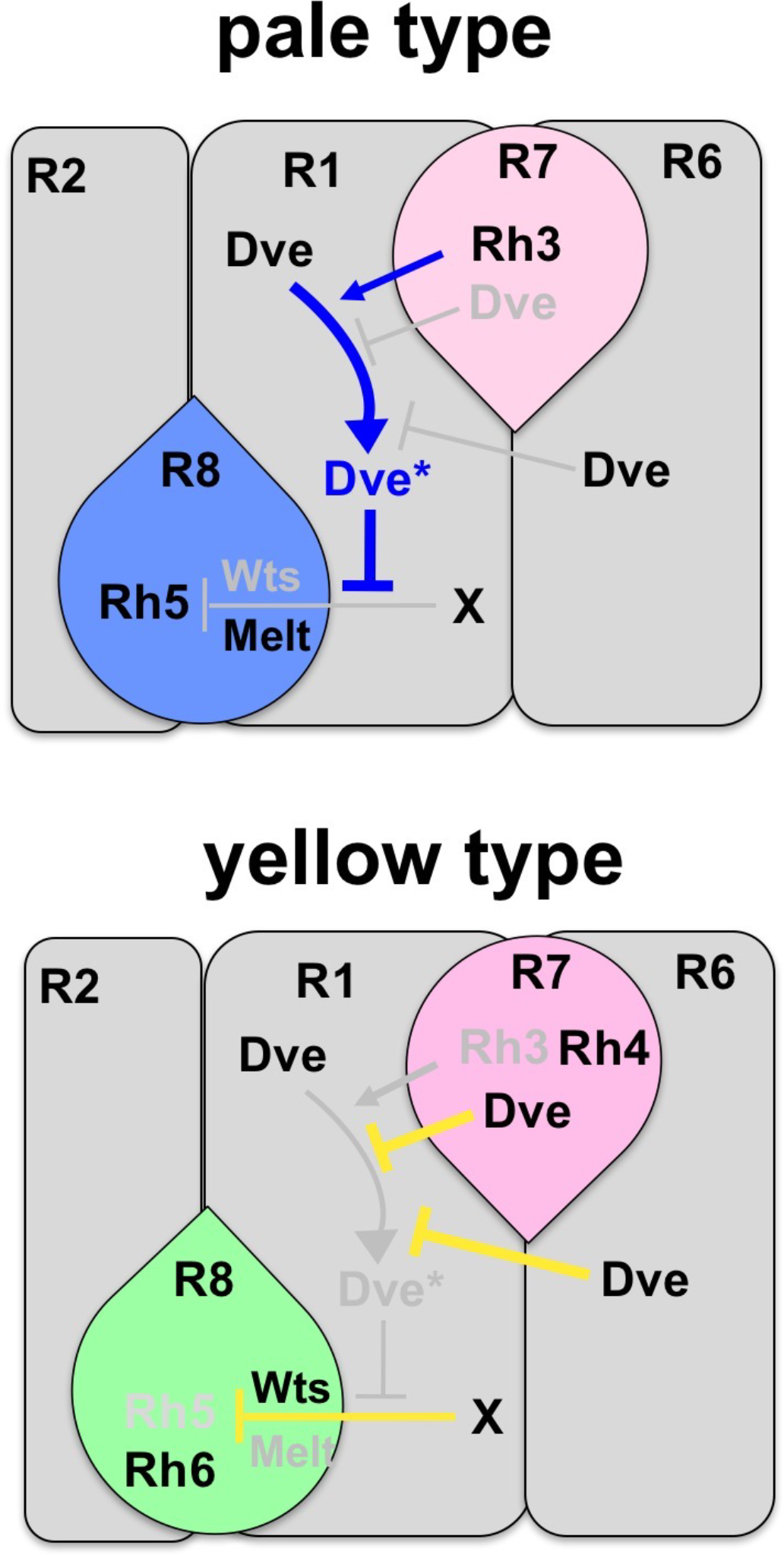
A model of Rh5-instructive signal transmission. Rh5 expression in the R8 is repressed by the Hippo signaling (Wts/Melt) through unknown signal “X”. In pale-type ommatidia, Rh3-expressing R7 sends a signal to the adjacent R1. The activated form of Dve (Dve*) in the R1 represses the signal “X” and induces Rh5 expression through a relief of repression mechanism. In yellow-type ommatidia, Spineless (Ss) induces Rh4 and Dve expression in the R7, and Dve represses Rh3 expression (Johnston et al., 2011). Dve activities in the R7 and R6 repress the activation of Dve in the R1 and lead to a default state of Rh6 expression.

In mouse retinas, thyroid hormone (TH) and TH receptor β2 is required to activate M-opsin and to repress S-opsin (Glaschke et al., 2011; Ng et al., 2001; Roberts et al., 2006). These reports suggest that extrinsic signals are required for spatial distribution and their subtype specification of cone cells. Moreover, our results raise an intriguing possibility that local intercellular signaling between rod and cone cells is also important for subtype specification. Thus, further characterization of *Drosophila* color vision will provide insights into the mechanism of functional specification during retinal development.

## Materials and Methods

### *Drosophila* strains

All flies were reared on a standard yeast and cornmeal-based diet at 25°C. Oregon-R (OR) flies were used as wild-type controls. *dve*^*1*^ is a severe loss of function allele that has no *dve-A* and a very weak *dve-B* activity in the larval midgut (Nakagawa et al., 2011; Nakagoshi et al., 1998). *dve*^*L186*^ and *ss*^*D115*.*7*^ are null alleles (Duncan et al., 1998; Terriente et al., 2008). FLP lines that express FLP in the eye-antennal disc (*ey-flp2*, BL5580) and that in R1, R6, and R7 (*GMR-flp*, BL42735) were used. *GMR-wIR* (gift from R. Carthew) was used to induce *white* RNAi in adult eyes (Lee and Carthew, 2003).

### Mosaic analyses

Mutant mosaic clones were induced by the FRT- and FLP-mediated recombination system (Xu and Rubin, 1993) as following genotypes:

*yw ey-flp2/Y; FRT42D dve*^*1*^*/FRT42D w*^*+*^ *M(2)53*^*1*^,

*yw ey-flp2/Y; FRT42D dve*^*L186*^*/FRT42D w*^*+*^ *M(2)53*^*1*^,

*yw ey-flp2/Y; FRT42D dve*^*1*^*/+; FRT82B ss*^*D115*.*7*^*/FRT82B w*^*+*^ *l(3)cl-R3*

*yw ey-flp2/Y; FRT42D dve*^*1*^*/FRT42 GMR-hid; FRT82B ss*^*D115*.*7*^*/FRT82B w*^*+*^ *l(3)cl-R3*

*yw eyflp2/GMR-wIR; FRT42D dve*^*L186*^*/+; FRT82B ss*^*D115*.*7*^*/FRT82B GMR-hid l(3)CL-R1*

*yw eyflp2/GMR-wIR; FRT42D dve*^*L186*^*/FRT42 GMR-hid; FRT82B ss*^*D115*.*7*^*/FRT82B GMR-hid l(3)CL-R1*

Mutant mosaic clones labeled with GFP were induced by the MARCM system (Lee and Luo, 1999) as following genotypes:

*GMR-flp yw/UAS-ActGFP; FRTG13 UAS-mCD8-GFP dve*^*1*^*/FRTG13 tub-GAL80; tub-GAL4/GMR-wIR*

*GMR-flp yw/UAS-ActGFP; FRTG13 UAS-mCD8-GFP dve*^*L186*^*/FRTG13 tub-GAL80; tub-GAL4/GMR-wIR*

### Immunohistochemistry

Adult compound eyes were dissected in phosphate-buffered saline (PBS), fixed with 4% formaldehyde/PBS-0.3% Triton X-100 for 15 min, and washed several times with PBS-0.3% Triton X-100. The following primary antibodies were used: rabbit anti-Dve (1:2000) (Nakagoshi et al., 1998), mouse anti-Rh3 (2B1, gift from S. Britt, 1:20), rabbit anti-Rh4 (gift from C. Zucker, 1:200), mouse anti-Rh5 (7F1, gift from S. Britt, 1:200), rabbit anti-Rh6 (gift from C. Desplan, 1:2000). FITC-, Cy3- or Cy5-conjugated secondary antibodies (Jackson Immunoresearch) were used for detection. Phalloidin-TRITC (Sigma) was used to stain actin fibers of rhabdomere. Confocal images of 0.2–1.22 μm sections were obtained with a confocal microscope (Olympus FV1200) and were processed using the Fluoview (Olympus) and Photoshop software (Adobe).

### Statistical analysis

The significance of differences between the control and test progenies was analyzed using Prism6 (GraphPad Software). A chi-square test was applied to compare the frequency distribution of Rhodopsin pairing (ommatidial subtypes). Fisher’s exact test was applied to compare the frequency distribution of Rhodopsin expression in R8. The levels of significance are indicated by asterisks: \**p* < 0.05, \*\**p* < 0.01, \*\*\**p* < 0.001.

## Acknowledgments

We are grateful to Steven Britt, Charles Zucker, Claude Desplan for antibodies, Richard Carthew for *GMR-wIR*, the Bloomington Drosophila Stock Center, the Vienna Drosophila Resource Center (VDRC), the Drosophila Genomics and Genetic Resources (DGGR, Kyoto Stock Center), and the National Institute of Genetics (NIG) for fly strains. We thank the Developmental Studies Hybridoma Bank (DSHB) for antibodies. We thank Claude Desplan and Robert Johnston for helpful discussion.

## References

Anderson, C., Reiss, I., Zhou, C., Cho, A., Siddiqi, H., Mormann, B., Avelis, C. M., Deford, P., Bergland, A., Roberts, E., et al. (2017). Natural variation in stochastic photoreceptor specification and color preference in Drosophila. Elife 6, e29593.

Birkholz, D. A., Chou, W. H., Phistry, M. M. and Britt, S. G. (2009a). Disruption of photoreceptor cell patterning in the Drosophila Scutoid mutant. Fly (Austin) 3, 253–262.

Birkholz, D. A., Chou, W. H., Phistry, M. M. and Britt, S. G. (2009b). rhomboid mediates specification of blue- and green-sensitive R8 photoreceptor cells in Drosophila. J Neurosci 29, 2666–2675.

Bumsted O’Brien, K. M., Cheng, H., Jiang, Y., Schulte, D., Swaroop, A. and Hendrickson, A. E. (2004). Expression of photoreceptor-specific nuclear receptor NR2E3 in rod photoreceptors of fetal human retina. Invest. Ophthalmol. Vis. Sci. 45, 2807–2812.

Cheng, H., Khan, N. W., Roger, J. E. and Swaroop, A. (2011). Excess cones in the retinal degeneration rd7 mouse, caused by the loss of function of orphan nuclear receptor Nr2e3, originate from early-born photoreceptor precursors. Hum. Mol. Genet. 20, 4102–4115.

Cheng, H., Khanna, H., Oh, E. C., Hicks, D., Mitton, K. P. and Swaroop, A. (2004). Photoreceptor-specific nuclear receptor NR2E3 functions as a transcriptional activator in rod photoreceptors. Hum. Mol. Genet. 13, 1563–1575.

Chou, W.-H., Huber, A., Bentrop, J., Schulz, S., Schwab, K., Chadwell, L. V., Paulsen, R. and Britt, S. G. (1999). Patterning of the R7 and R8 photoreceptor cells of Drosophila: evidence for induced and default cell-fate specification. Development 126, 607–616.

Cook, T., Pichaud, F., Sonneville, R., Papatsenko, D. and Desplan, C. (2003). Distinction between color photoreceptor cell fates is controlled by Prospero in Drosophila. Dev. Cell 4, 853–864.

Duncan, D. M., Burgess, E. A. and Duncan, I. (1998). Control of distal antennal identity and tarsal development in Drosophila by spineless-aristapedia, a homolog of the mammalian dioxin receptor. Genes Dev. 12, 1290–1303.

Glaschke, A., Weiland, J., Del Turco, D., Steiner, M., Peichl, L. and Glosmann, M. (2011). Thyroid hormone controls cone opsin expression in the retina of adult rodents. J. Neurosci. 31, 4844–4851.

Haider, N. B., Jacobson, S. G., Cideciyan, A. V., Swiderski, R., Streb, L. M., Searby, C., Beck, G., Hockey, R., Hanna, D. B., Gorman, S., et al. (2000). Mutation of a nuclear receptor gene, NR2E3, causes enhanced S cone syndrome, a disorder of retinal cell fate. Nat. Genet. 24, 127–131.

Johnston, R. J., Jr., Otake, Y., Sood, P., Vogt, N., Behnia, R., Vasiliauskas, D., McDonald, E., Xie, B., Koenig, S., Wolf, R., et al. (2011). Interlocked feedforward loops control cell-type-specific Rhodopsin expression in the Drosophila eye. Cell 145, 956–968.

Jukam, D. and Desplan, C. (2011). Binary regulation of Hippo pathway by Merlin/NF2, Kibra, Lgl, and Melted specifies and maintains postmitotic neuronal fate. Dev Cell 21, 874–887.

Jukam, D., Xie, B., Rister, J., Terrell, D., Charlton-Perkins, M., Pistillo, D., Gebelein, B., Desplan, C. and Cook, T. (2013). Opposite feedbacks in the Hippo pathway for growth control and neural fate. Science 342, 1238016.

Lee, T. and Luo, L. (1999). Mosaic analysis with a repressible cell marker for studies of gene function in neuronal morphogenesis. Neuron 22, 451–461.

Lee, Y. S. and Carthew, R. W. (2003). Making a better RNAi vector for Drosophila: use of intron spacers. Methods 30, 322–329.

Mikeladze-Dvali, T., Wernet, M. F., Pistillo, D., Mazzoni, E. O., Teleman, A. A., Chen, Y. W., Cohen, S. and Desplan, C. (2005). The growth regulators warts/lats and melted interact in a bistable loop to specify opposite fates in Drosophila R8 photoreceptors. Cell 122, 775–787.

Mollereau, B. and Domingos, P. M. (2005). Photoreceptor differentiation in Drosophila: from immature neurons to functional photoreceptors. Dev. Dyn. 232, 585–592.

Mollereau, B., Dominguez, M., Webel, R., Colley, N. J., Keung, B., de Celis, J. F. and Desplan, C. (2001). Two-step process for photoreceptor formation in Drosophila. Nature 412, 911–913.

Nakagawa, Y., Fujiwara-Fukuta, S., Yorimitsu, T., Tanaka, S., Minami, R., Shimooka, L. and Nakagoshi, H. (2011). Spatial and temporal requirement of Defective proventriculus activity during Drosophila midgut development. Mech. Dev. 128, 258–267.

Nakagoshi, H., Hoshi, M., Nabeshima, Y. and Matsuzaki, F. (1998). A novel homeobox gene mediates the Dpp signal to establish functional specificity within target cells. Genes Dev. 12, 2724–2734.

Nathans, J. (1999). The evolution and physiology of human color vision: insights from molecular genetic studies of visual pigments. Neuron 24, 299–312.

Ng, L., Hurley, J. B., Dierks, B., Srinivas, M., Salto, C., Vennstrom, B., Reh, T. A. and Forrest, D. (2001). A thyroid hormone receptor that is required for the development of green cone photoreceptors. Nat. Genet. 27, 94–98.

Peng, G.-H., Ahmad, O., Ahmad, F., Liu, J. and Chen, S. (2005). The photoreceptor-specific nuclear receptor Nr2e3 interacts with Crx and exerts opposing effects on the transcription of rod versus cone genes. Hum. Mol. Genet. 14, 747–764.

Roberts, M. R., Srinivas, M., Forrest, D., Morreale de Escobar, G. and Reh, T. A. (2006). Making the gradient: thyroid hormone regulates cone opsin expression in the developing mouse retina. Proc. Natl. Acad. Sci. U. S. A. 103, 6218–6223.

Tahayato, A., Sonneville, R., Pichaud, F., Wernet, M. F., Papatsenko, D., Beaufils, P., Cook, T. and Desplan, C. (2003). Otd/Crx, a dual regulator for the specification of ommatidia subtypes in the Drosophila retina. Dev. Cell 5, 391–402.

Tan, H., Fulton, R. E., Chou, W. H., Birkholz, D. A., Mannino, M. P., Yamaguchi, D. M., Aldrich, J. C., Jacobsen, T. L. and Britt, S. G. (2020). Drosophila R8 photoreceptor cell subtype specification requires hibris. PLoS One 15, e0240451.

Terriente, J., Perea, D., Suzanne, M. and Diaz-Benjumea, F. J. (2008). The Drosophila gene zfh2 is required to establish proximal-distal domains in the wing disc. Dev. Biol. 320, 102–112.

Thanawala, S. U., Rister, J., Goldberg, G. W., Zuskov, A., Olesnicky, E. C., Flowers, J. M., Jukam, D., Purugganan, M. D., Gavis, E. R., Desplan, C., et al. (2013). Regional modulation of a stochastically expressed factor determines photoreceptor subtypes in the Drosophila retina. Dev Cell 25, 93–105.

Wells, B. S., Pistillo, D., Barnhart, E. and Desplan, C. (2017). Parallel Activin and BMP signaling coordinates R7/R8 photoreceptor subtype pairing in the stochastic Drosophila retina. Elife 6, e25301.

Wernet, M. F. and Desplan, C. (2004). Building a retinal mosaic: cell-fate decision in the fly eye. Trends Cell Biol. 14, 576–584.

Wernet, M. F., Mazzoni, E. O., Celik, A., Duncan, D. M., Duncan, I. and Desplan, C. (2006). Stochastic spineless expression creates the retinal mosaic for colour vision. Nature 440, 174–180.

Xu, T. and Rubin, G. M. (1993). Analysis of genetic mosaics in developing and adult Drosophila tissues. Development 117, 1223–1237.

